# Antimicrobial-producing Bacteria from Fish Epidermal Mucus Alter the Fish Epidermal Bacterial Flora and Host Resistance to Infection

**DOI:** 10.1101/2025.07.17.665321

**Authors:** Hajime Nakatani, Naoto Suetake, Katsutoshi Hori

## Abstract

The emergence of antimicrobial-resistant bacteria in aquaculture has raised the need for alternative strategies to control fish infections. Antimicrobial-producing bacteria have been explored as probiotics or biocontrol agents, but their mechanisms of action and impact on host-associated microbiota remain poorly understood. Here, we identified *Pseudomonas mosselii* KH-ZF1, a bacterium isolated from fish epidermal mucus, as a producer of antimicrobial substances. When applied to zebrafish, KH-ZF1 transiently adhered to the epidermal mucus and altered the composition of the skin microbiota. Under an appropriate administration condition, KH-ZF1 treatment significantly improved survival in zebrafish infected with *Yersinia ruckeri,* and suppressed pathogen growth on the skin surface. However, in the absence of KH-ZF1 or inappropriate conditions, *Y. ruckeri* dominated the epidermal bacterial community. The antimicrobial compound produced by KH-ZF1 was identified as Fluviol C (FluC), a pigmented metabolite previously reported from *Pseudomonas fluorescens*. FluC inhibited the growth of multiple fish pathogens at low concentrations, but exhibited toxicity to zebrafish even below its minimum inhibitory concentration. Intriguingly, FluC at sub-inhibitory levels induced bacterial substitution in the epidermal microbiota, mimicking the effects of KH-ZF1. These findings demonstrate that KH-ZF1 alters host resistance to infection by promoting bacterial substitution on the fish skin by producing FluC. Our study highlights a microbiota-mediated mechanism by which antimicrobial-producing bacteria can control infection through fish epidermis, offering a potential alternative to traditional antibiotics in aquaculture.

**Importance:** We demonstrated for the first time that bacteria producing antibacterial substances, isolated from fish skin mucus, can inhibit percutaneous infections in aquatic environments. These bacteria effectively altered the skin mucus bacterial flora and suppressed pathogen growth. Fish skin acts as a barrier against infections, with its microorganisms being considered to play a crucial role in prevention. Our study highlighted the potential use of these specific microorganisms in the fish skin mucus as a novel fish disease control strategy. By targeting fish skin mucus bacteria that produce antimicrobial substances, we could develop a new approach to managing diseases in aquaculture like probiotics for fish skin. This research underscores the importance of studying fish epidermal microorganisms for innovative disease management.

## Introduction

Disease control is a critical challenge for improving productivity and sustainability in aquaculture. Infectious diseases are a leading cause of economic loss in fish farming, and antimicrobial agents have been widely adopted due to their ease of use and broad-spectrum efficacy. However, the increasing use of these agents has contributed to the emergence and spread of antimicrobial-resistant bacteria, which not only reduces treatment effectiveness but also poses a significant risk to human health by complicating the treatment of infectious diseases (1–3). This global concern has driven the search for alternative and more sustainable disease control strategies.

One promising approach is the use of beneficial microorganisms for biological control, which has been extensively explored in agriculture (4, 5). In aquaculture, livestock systems and medical care, probiotics, live microorganisms that confer health benefits to the host, have been administered orally to improve disease resistance and overall health. Various strains have been reported to inhibit pathogen colonization, enhance nutrient absorption, modulate immune responses, and competitively exclude harmful microbes (6–12). Among these functions, the ability to suppress pathogenic bacteria through the production of antimicrobial substances is often considered a key screening criterion for beneficial microorganisms (13–17). However, few studies have examined how these antimicrobial substances from the microorganisms affect the microbial community, including both pathogenic and non-pathogenic microorganisms. Understanding these effects is essential for effective infection prevention (8, 18).

Like the intestinal tract, the fish epidermis is covered by a mucus layer that harbors a complex and dynamic microbial community. This skin-associated microbiota functions as a critical barrier against environmental pathogens, especially in aquatic environments where the skin is in constant contact with waterborne microbes (19–21). However, the application of antimicrobial-producing microorganisms like probiotics has focused primarily on the intestinal microbiota, and relatively few studies have addressed whether these microorganisms positively affect the microbial communities on fish epidermal mucus.

Recent studies, including our own, have suggested that disturbances in the epidermal microbial community, whether due to environmental stress, antimicrobial agents, or shifts in microbial composition, can alter disease susceptibility (22, 23). In zebrafish (*Danio rerio*), for example, reduced water temperatures have been shown to facilitate percutaneous infections by fish pathogens, accompanied by simultaneous changes in the skin microbiota (23). We have also observed that increased abundance of antimicrobial- producing bacteria, as well as exposure to antibiotics, can significantly alter the composition of the epidermal bacterial flora, sometimes leading to the dominance of pathogenic or opportunistic bacteria (22, 23). These findings suggest that antimicrobial- producing bacteria may exert similar impacts to fish skin microflora as antibiotics, but the underlying mechanisms remain poorly understood.

A more detailed understanding of how antimicrobial-producing bacteria interact with host-associated microbial communities, particularly on the skin, is essential for developing disease control strategies for aquaculture. While much of the focus in such research has been on gut-associated microbes, the fish skin represents a promising but underexplored site for microbial intervention.

In this study, we focused on the epidermal mucus microbiota of zebrafish and identified *Pseudomonas mosselii* strain KH-ZF1, an antimicrobial-producing bacterium isolated from the epidermal mucus. We evaluated whether the addition of this strain to aquarium water could prevent percutaneous infection by the fish pathogen *Yersinia ruckeri*, and analyzed its effects on the composition of the epidermal bacterial community. Moreover, we identified the antimicrobial compound produced by KH-ZF1 and examined its specific effects on pathogen growth and bacterial flora on zebrafish skin. This study aims to deepen our understanding of how antimicrobial-producing bacterial inputs influence the fish skin microbiota and host infection outcomes, with implications for skin microbiota- targeted disease prevention in aquaculture.

## Results

### Isolation of Bacteria from Fish Epidermal Mucus to Inhibit the Growth of Fish Pathogens

To identify bacteria capable of inhibiting fish pathogens, we screened isolates from the epidermal mucus of zebrafish. Using the cross-streak method, six clones (C6, KH-ZF1, N5, N10, C5, and m4-T2) were found to exhibit antagonistic activity against various fish pathogens (Table 1 and Fig. 1A). Among them, clone KH-ZF1 and C6 inhibited multiple pathogens. Phylogenetic analysis based on 16S rDNA sequences identified KH-ZF1 and C6 as closely related to *Pseudomonas mosselii* and *Brevibacterium casei*, respectively (Fig. S1). Amplicon sequence variant (ASV)-based bacterial community analysis of zebrafish epidermal mucus, the isolation source of strains C6 and KH-ZF1, and rearing water was performed, and the relative abundances of the top 30 bacterial genera were shown (Fig. 2). The genus *Pseudomonas* accounted for 6–10% of the community (Fig. 2A). Notably, *Brevibacterium*, from which strain C6 was isolated, was not detected in the community. Phylogenetic analysis of ASVs assigned to the genus *Pseudomonas* revealed that a majority were affiliated with *Pseudomonas mosselii* (Fig. 2B). Among the *Pseudomonas* species, *P. mosselii* was second predominant species in the epidermal mucus, comprising 1.43% of total ASVs. Other frequently detected species included *P. lutea* (2.88%) and *P. parafulva* (1.02%)(Fig. 2C). Accordingly, KH-ZF1 was selected for further experiments as an antimicrobial-producing bacterium existing in fish epidermal mucus. Additional tests confirmed KH-ZF1’s broad antimicrobial activity against fish pathogens except for *Acinetobacter* sp. (Fig. S2).

**Fig. 1.**
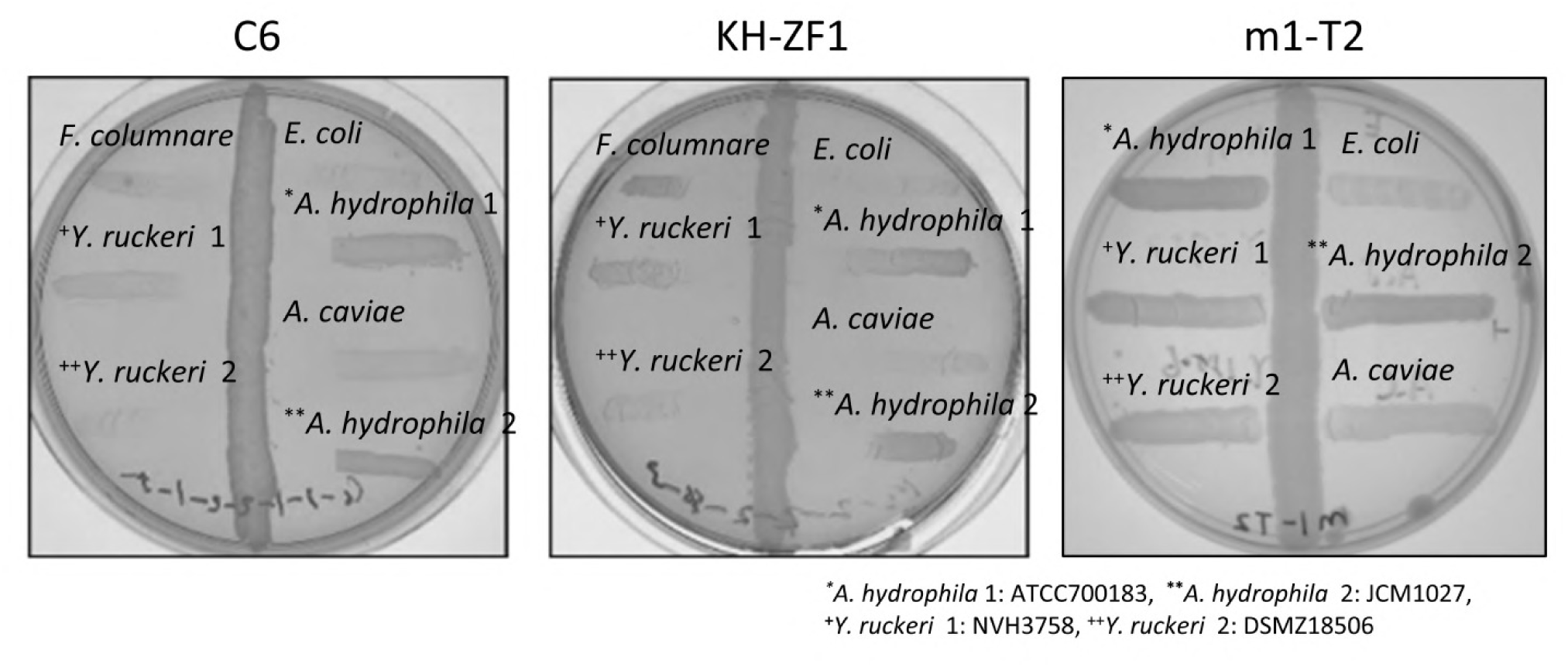
KH-ZF1 and C6 strains inhibit the growth of fish pathogens. Antimicrobial activity of C6, KH-ZF1, and m1-T2 (negative control) clones against fish pathogens were examined by cross-streak method.

**Fig. 2.**
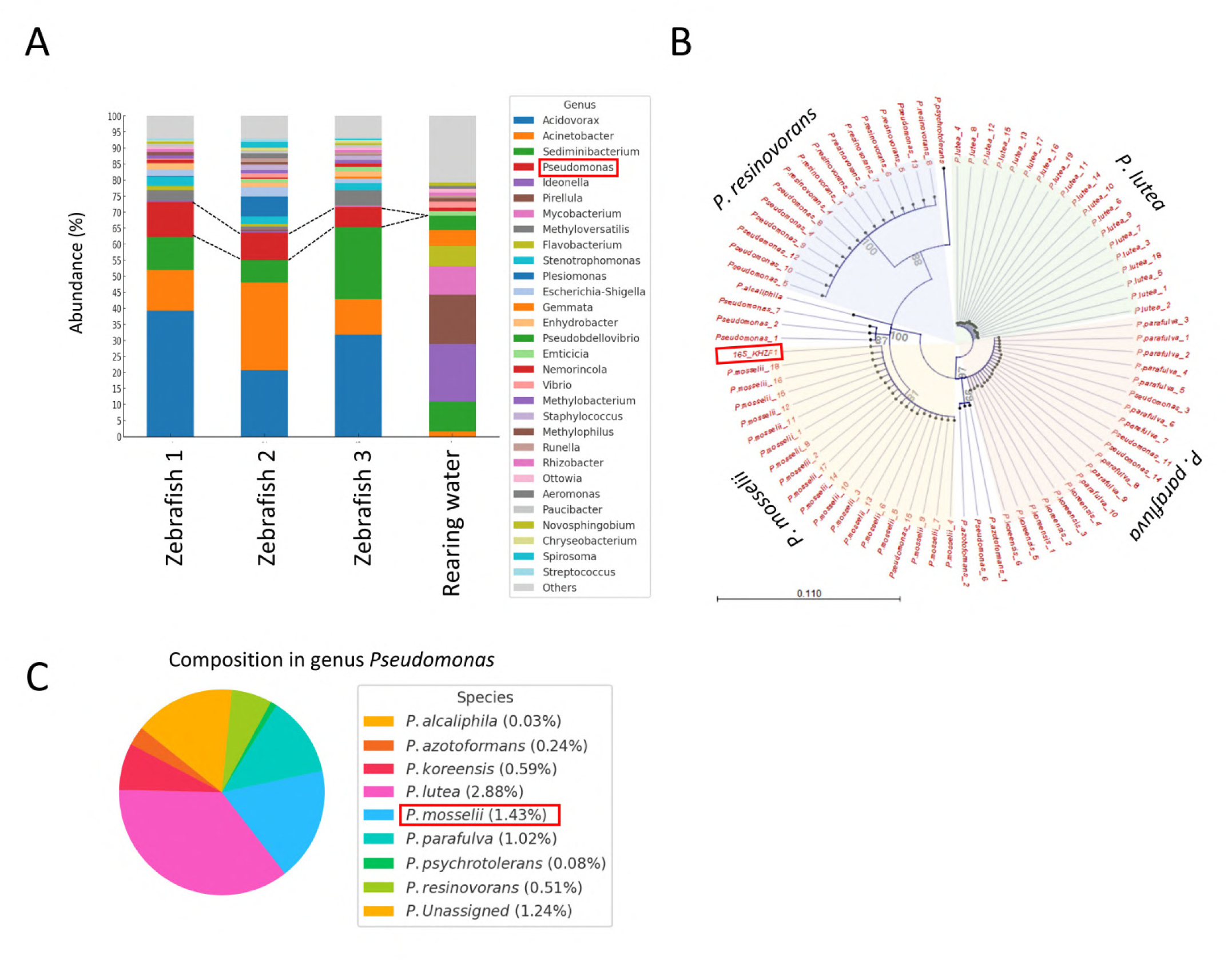
Bacterial community composition of the isolation sources for KH-ZF1 and phylogenetic analysis of the genus *Pseudomonas* in zebrafish epidermal mucus. (A) Bacterial community composition in the isolation sources (zebrafish epidermal mucus and rearing water) for strains C6 and KH-ZF1 was analyzed based on ASV clustering. The relative abundances of the top 30 genera, including *Pseudomonas* (6–10%), are shown. *Brevibacterium* (strain C6) was not detected in this analysis. (B) Phylogenetic analysis of ASVs assigned to the genus *Pseudomonas*. KH-ZF1 was located within the clade of *Pseudomonas mosselii*. (C) Relative abundance of *Pseudomonas mosselii* among major *Pseudomonas* species present in the zebrafish epidermal mucus. *P. mosselii* accounted for 1.43% of the total bacterial community. Other dominant *Pseudomonas* species included *P. lutea* (2.88%) and *P. parafulva* (1.02%).

**Table 1.** The zebrafish epidermal bacteria with growth-inhibitory effect against fish pathogens.

| Pathoges<br>Clones | *Aeromonas caviae*<br>JCM1043 | *Aeromonas hydrophila*<br>JCM1027 | *Flavobacterium columnare*<br>JCM21327 | *Yersinia ruckeri*<br>NVH3758 |
| --- | --- | --- | --- | --- |
| C6 | + | + | + | + |
| KH-ZF1 | + | + | + | + |
| N5 | - | - | + | - |
| N10 | - | - | + | - |
| C5 | - | - | + | - |
| m4-T2 | + | - | n.d. | - |

+ : Positive (Clear zoon 10 mm >) - : Negative n.d. : Not determine

### Microbial Substitution in the Epidermal Mucus Bacterial Flora by Administration of KH-ZF1

We assessed whether KH-ZF1 could establish on the fish epidermis and alter the resident microbial community. KH-ZF1 was introduced into rearing water under conditions favorable to *Yersinia ruckeri* percutaneous infection (23). Zebrafish were maintained at 20°C with epidermal injuries, and KH-ZF1::*mChery* (genetically tagged with *mCherry* and kanamycin resistance gene) was used for following experiments. For the initial step of CFU measurement of KH-ZF1 on fish epidermal mucus, mucus samples were collected from zebrafish treated with KH-ZF1::*mCherry* and from untreated controls. These samples were incubated on a selective medium specific for KH-ZF1::*mCherry*. As shown in Fig. S3, fluorescent colonies were obtained only from the KH-ZF1::*mCherry*-treated samples, confirming the selective detection of KH-ZF1::*mCherry* on the medium.

On the first day after the initial dose, the number of KH-ZF1 cells on the zebrafish epidermal mucus was approximately 10^6^-10^7^ CFU per fish one day after the initial dose and declined to 10^4^ CFU per fish by days two and three. A second dose at day one hardly affected the number of KH-ZF1 cells on the epidermis, suggesting 10⁶-10⁷ CFU per fish is maximum number of KH-ZF1 cells on the epidermis. A third dose at day three recovered the number of KH-ZF1 cells to 10^6^ CFU per fish on the first day after the third dose. By the seventh day after the initial dose, the number of KH-ZF1 cells remained at approximately 10^5^ CFU per fish when the third dose was done at day three. In contrast, without third dose, the number of KH-ZF1 cells decreased to 10^3^ CFU per fish at day seven (Fig. 3A). Observations of KH-ZF1 cells on the epidermis after the second dose revealed that the KH-ZF1 aggregates were sparsely distributed on the surface of the fish, with notable localization at the wounded sites on the epidermis (Fig. 3B).

**Fig. 3.**
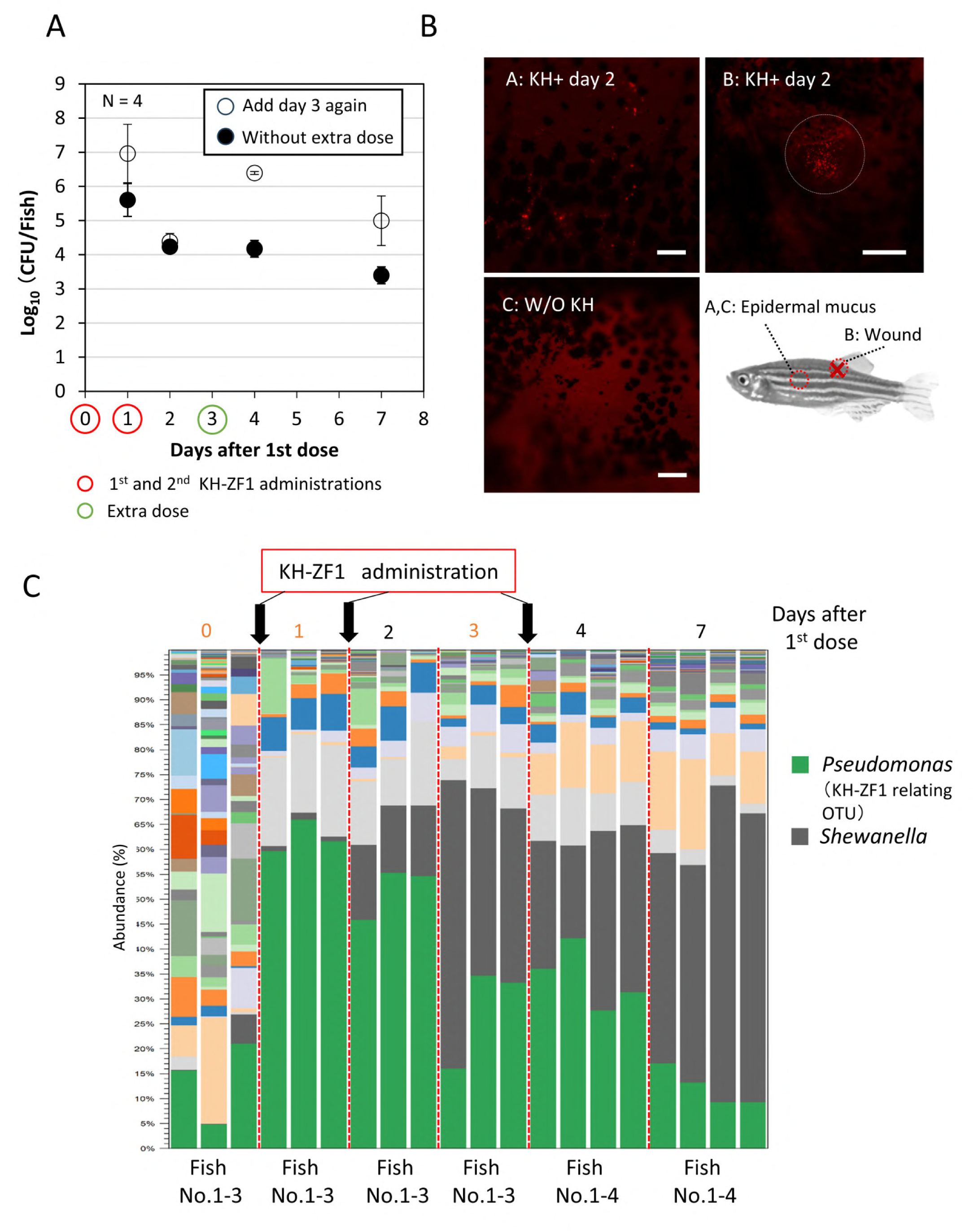
KH-ZF1 administration increases the number of the cells and relative abundance in the epidermal bacterial flora, and promotes microbial substitution. (A) Colony forming unit (CFU) of KH-ZF1 was measured at several days after administrations to the injured zebrafish. (B) KH-ZF1 cells producing mCherry were administrated and the cells on the epidermis were observed at 2 days after the first dose under fluorescence microscope. The KH-ZF1 cells producing mCherry were observed as small red spot. (C) The bacterial composition on the epidermis was analyzed before KH-ZF1 administration and at several days after administration of KH-ZF1. OTUs assigned KH-ZF1-related *Pseudomonas* and *Shewanella* were shown.

To analyze the effect of successive KH-ZF1 administrations on the epidermal microflora, the bacterial flora analysis during KH-ZF1 administration was carried out. The proportion of the KH-ZF1-related OTUs in the bacterial community increased significantly at day one. However, the occupancy rate of these OTUs gradually decreased, even after the second and third doses were administered (Fig. 3C). Notably, the proportion of *Shewanella* OTUs expanded substantially from 1-10% to 44-64% by day seven by KH- ZF1 administration (Fig. 3C). In the control group without KH-ZF1 administration, injury and a reduction in water temperature to 20°C transiently affected the epidermal bacterial community. Notably, genera such as *Shewanella* and *Aeromonas*, which are capable of growing at lower temperatures, temporarily became dominant in the skin mucus. However, the dominance of these OTUs diminished over the subsequent days (Fig. S4). In contrast to the KH-ZF1-treated group, OTUs related to KH-ZF1 did not become dominant at any point during the experiment (Fig. 3C and Fig. S4). These findings indicate that the observed shifts in the bacterial community—including the increased abundance of KH-ZF1-related taxa and broader compositional changes—were induced by KH-ZF1 administration.

Similar microbial substitutions were observed in the gills and intestinal content (Fig. S5), suggesting that KH-ZF1 can broadly modulate mucosal microbiota, including that of the fish epidermis.

### Protection of Percutaneous Infection of *Yersinia ruckeri* by Multiple Administrations of KH-ZF1

To evaluate the protective effect of KH-ZF1 in rearing water against percutaneous infection, zebrafish were challenged with *Y. ruckeri* as previous study (23) and survival was monitored under various dosing regimens. Fish were subjected to injury and subsequently transferred to flasks with adjusted water temperature. Following this, they were challenged with *Yersinia ruckeri*. These experiments were repeated three to four times, and datasets necessary for constructing Kaplan–Meier survival curves were obtained (Fig. S6).

Single-dose treatments showed no effect on the survival rate at eight days after pathogen challenge when compared to the pathogen challenge only group (Fig. 4A). Single-dose of KH-ZF1 before the pathogen challenge (KHx1 before day0), and one day after the pathogen challenge (KHx1 day1) appeared to prolong the survival of the fish. However, immediate administration after the pathogen challenge (KHx1 day0) seemed to decrease the survival rate at day eight (Fig. 4A).

**Fig. 4.**
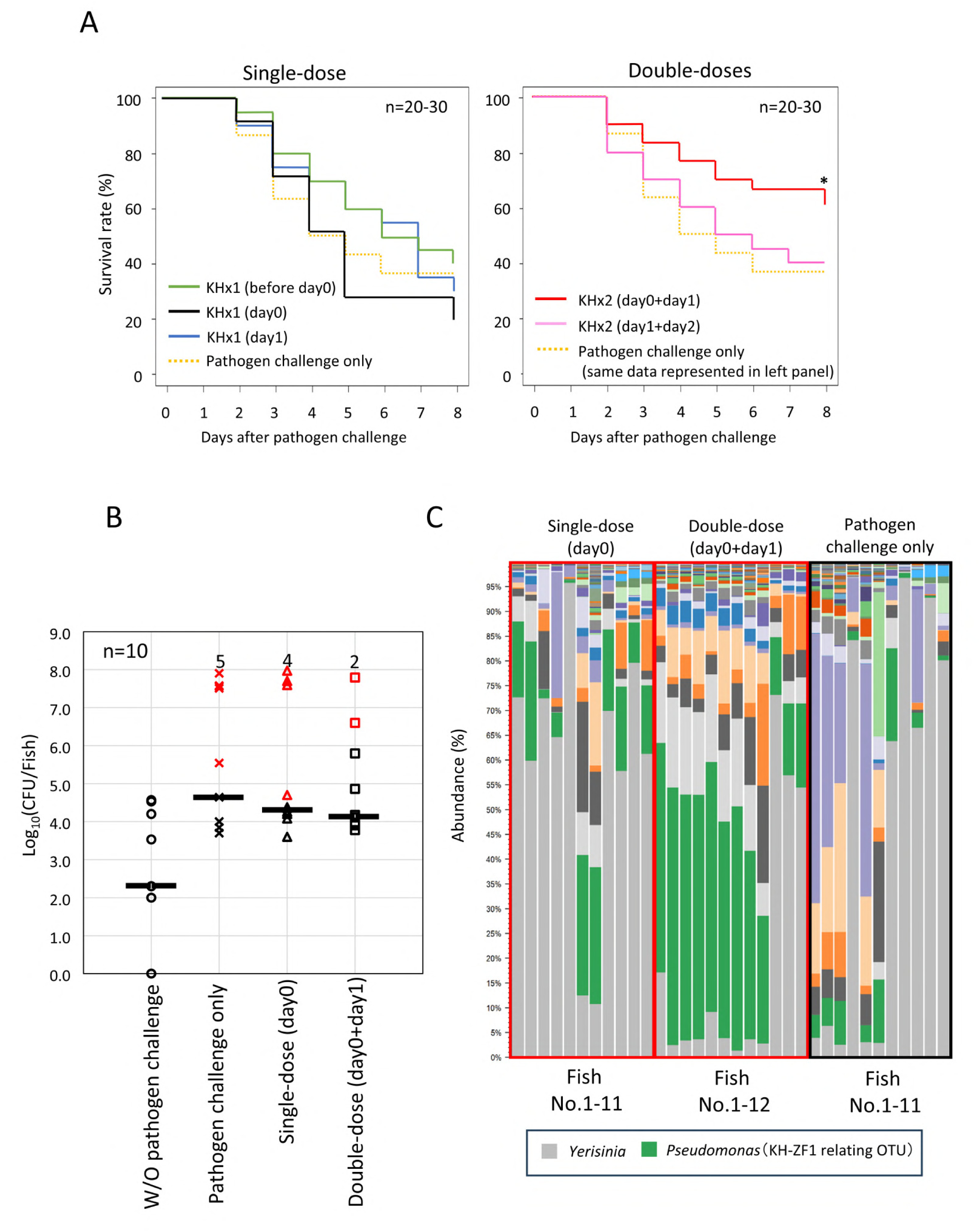
KH-ZF1 administration prevent percutaneous infection of *Yersinia ruckeri* by influencing the growth of pathogen on the epidermis and inducing microbial substitution. (A) Transition of the survival rate of zebrafish after pathogen challenge at day 0 and single-dose or double-dose KH-ZF1 administration. Asterisk (*) represents significant increase of survival rate against ‘pathogen challenge only’ group (Log-rank test p<0.05). (B) CFU *of Y. ruckeri* on the epidermis of the fish without pathogen challenge, with pathogen challenge, and with KH-ZF1 administration after pathogen challenge (Single-dose (day0) and Double-dose (day0+day1)) was measured when the survival rate of ‘pathogen challenge only’ group became under 50%. The red-colored data points represent the values of dead fish. The number of dead fish in each group are expressed by red text. Mean values are expressed by bars. (C) The epidermal mucus bacterial flora of the fish with pathogen challenge and with KH-ZF1 administration after pathogen challenge (Single-dose (day0) and Double-dose (day0+day1)) was analyzed at three days after pathogen challenge.

In contrast, double-dose administration (immediately after and one day post-challenge: KHx2 (day0+day1)) significantly increased survival to 60-70% (Log-rank test *p*<0.05). The double-dose treatment only after pathogen challenge (KHx2 (day1+day2)) did not improve survival rate, indicating administration timing is critical. We next investigated the effect of KH-ZF1 administration on the growth of *Yersinia ruckeri* on the fish epidermis mucus. To measure the colony-forming units (CFU) of *Y. ruckeri* in the epidermal mucus, we used a strain of *Y. ruckeri* harboring *lacZ* and *Km^r^* genes in its genome (23). CFU measurements were taken when the survival rate of the pathogen challenge only group dropped below 50%. As shown in Fig. 4B, the CFU of *Y. ruckeri* in the epidermal mucus clearly increased following pathogen challenge compared to the control group (without pathogen challenge) regardless of the KH-ZF1 administration. In many fish without KH-ZF1 administration, the CFU of *Y. ruckeri* in the epidermal mucus reached 10^8^ CFU per fish. A single-dose treatment (day0) failed to suppress this increase in the number of *Y. ruckeri* in the epidermal mucus. However, a double-dose treatment (day0+day1) effectively suppresses the increase in the number of *Y. ruckeri* in the epidermal mucus of many fish. These results showed that multiple doses of KH-ZF1 at appropriate intervals inhibit the growth of *Y. ruckeri* in the epidermal mucus.

To further analyze the effect of microbial substitution induced by KH-ZF1 administration on the prevention of *Y. ruckeri* infection, we examined the epidermal mucus bacterial flora at three days after pathogen challenge during the infection experiments. Our previous study demonstrated that a challenge with *Y. ruckeri*, which also produces antimicrobial substances, promotes microbial substitution in the epidermal mucus, leading to the dominance of *Y. ruckeri* under conditions favorable for percutaneous infection (23). In this experiments, double-dose treatment (day0+day1) increased the abundance of KH-ZF1-related OTUs in the epidermal bacterial flora of many fish. In contrast, a single-dose (day0) or pathogen challenge only led to the occupation of *Y. ruckeri* in the bacterial flora of many fish (Fig. 4C). Consequently, multiple doses of KH-ZF1 at appropriate intervals increased the abundance of KH-ZF1 in the epidermal mucus through microbial substitution, thereby suppressing pathogen growth on the epidermis and improving survival rates. Conversely, inappropriate administration promoted the occupation of the pathogen in the bacterial flora, resulting in decreased survival rates.

### Identification of an antimicrobial substance produced by KH-ZF1

To isolate the antimicrobial substances produced by KH-ZF1 that are involved in inhibiting fish pathogens growth, we attempted to produce these products from liquid culture. First, we co-cultured KH-ZF1 with *Yersinia ruckeri* to determine when KH-ZF1 begins to produce antimicrobial substances. To verify production, we measured the CFU of *Y. ruckeri* at various time points during the culture. The CFU measurements indicated that the number of *Y. ruckeri* began to decrease around 12 hours of incubation when co- cultured with KH-ZF1 in NB medium, with a clear decrease observed after 24 hours (Fig. 5A), demonstrated that the antimicrobial activity of KH-ZF1 increased after 12 hours of incubation in liquid culture. Culture supernatants were then collected after 48 hours of incubation, and their antimicrobial activity was confirmed using the disk diffusion method. The supernatants from both the co-culture and single culture of KH-ZF1 contained antimicrobial substances that inhibited the growth of *Y. ruckeri* (Fig. 5B).

**Fig. 5.**
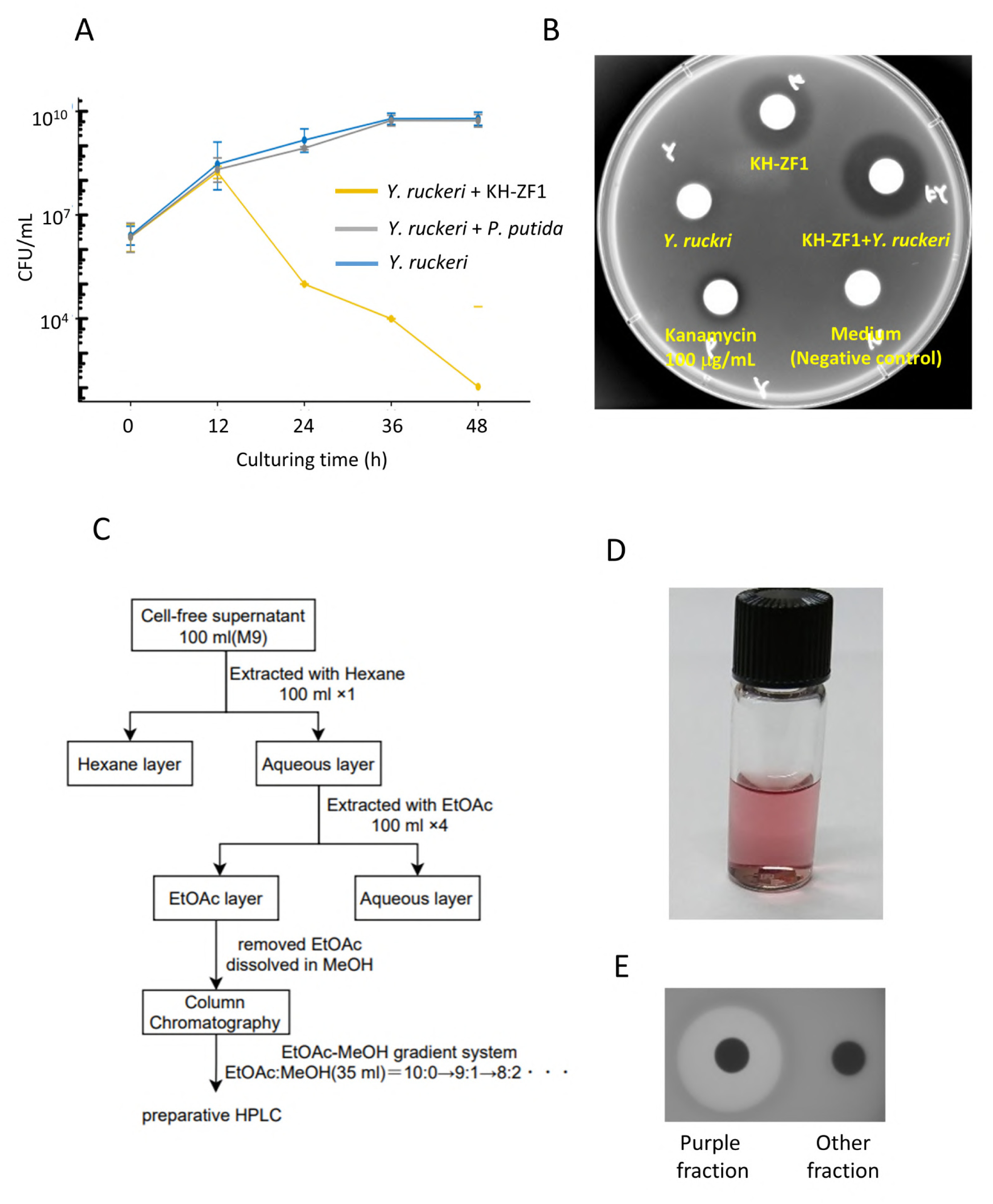
Isolation and purification of the antimicrobial substances produced by KH-ZF1. (A) CFU of *Y. ruckeri* in the co-culture with KH-ZF1 or Pseudomonas putida. (B) Detection of the antimicrobial activity in culture supernatants from single or co-culture of KH-ZF1 and/or *Y. ruckeri*. *Y.ruckeri* was used for a target pathogen. (C) Scheme for isolation and purification of the antimicrobial substances in the culture supernatant. (D) The fraction containing antimicrobial substances after HPLC. (E) Antimicrobial activity in the purple colored fraction and other fraction after HPLC were confirmed by disc diffusion method.

Next, we attempted to isolate and purify the antimicrobial substances from the culture supernatant. The extraction processes using various organic solvents were performed as shown in Fig. 5C. The fractions retaining antibacterial activity were separated by silica gel chromatography, and the active fraction was further purified by HPLC using a reversed-phase column. After separating the detected peaks at 220 nm by HPLC, we obtained a reddish-purple fraction containing the antimicrobial activity (Fig. 5D and 5E). This fraction was then assessed using LC-MS to determine its molecular mass. The total ion chromatogram showed a single chemical species in the fraction, and the MS of the ions generated from the chemical in the peak were measured as 166.0730 (+H), 188.0542 (+Na), and 204.0283 (+K), respectively (Fig. S7). These data indicate that the molecular mass of the antimicrobial substance is approximately 165.00-165.06. Using the Kazusa Molecular Formula Searcher (MFSearcher: https://webs2.kazusa-db.jp/mfsearcher/), we searched for compounds with compositions around MW 165.06 and identified possible candidates, including C_5_H_11_O_5_N, C_6_H_7_N_5_O, C_4_H_12_O_2_N_3_P, and C_9_H_12_NP.

The ^1^H-NMR spectra of this fraction showed two specific singlet peaks at 4.33 and 4.44 ppm (integral values: 19.64 and 20.23, respectively) corresponding to protons of methyl groups bound to alkynyl carbon, oxygen in ether groups, or nitrogen, and one specific singlet peak at 8.97 ppm (integral value: 6.00) corresponding to an aromatic proton (Fig. S8A). The ^13^C-NMR measurements revealed six specific peaks corresponding to six carbons in different environments, including two peaks around 40- 60 ppm from methyl carbons and four peaks around 140-160 ppm from aromatic carbons (Fig. S8B). Based on the NMR spectra, the most likely composition formula is C_6_H_7_N_5_O. Furthermore, the chemical shifts in the ^1^H-NMR and ^13^C-NMR suggested a chemical structure with a heteroaromatic compound containing methyl groups bound to nitrogen and oxygen. We predicted two possible chemical structures: known natural pyrazolotriazines named Fluviol C and Fluviol E (Fig. S8C).

To confirm the chemical structure of the antimicrobial substance, we crystallized it (Fig. 6A) and performed X-ray crystallography. The data obtained from crystallography are shown in Table S1, and the refined structure matched that of Fluviol C (IUPAC name: 3-Methoxy-7-methyl-7H-pyrazolo[4,3-E][1,2,4]triazine) (Fig. 6B). This substance is known as a pigment produced by *Pseudomonas fluorescens* var. *pseudoiodinum* (24) and was an identical chemical component recently reported as an antimicrobial substance produced by *Pseudomonas mosselii* 923, which inhibits plant pathogen infection (25).

**Fig. 6.**
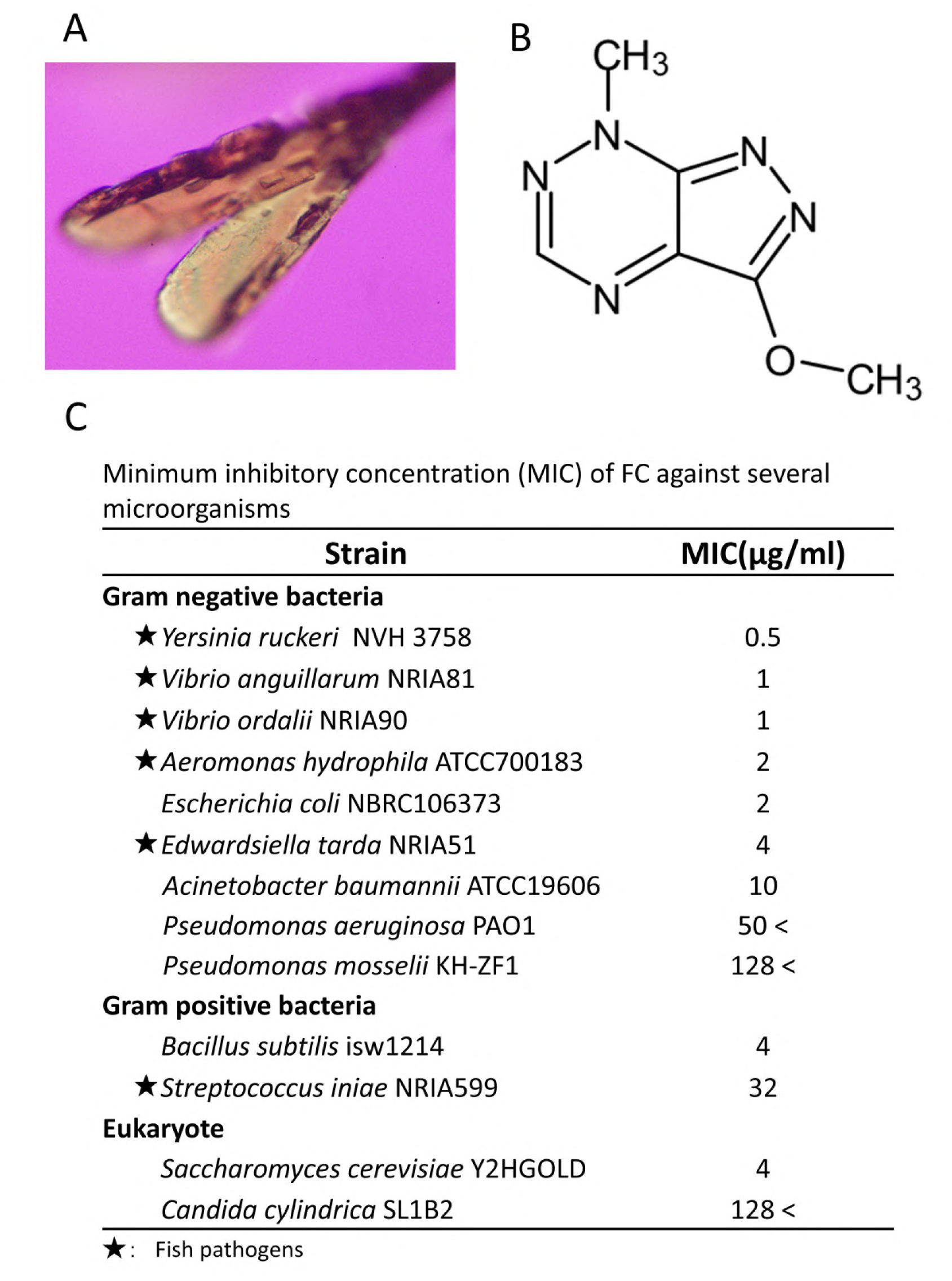
Fluviol C is an antimicrobial substance from KH-ZF1. (A) Crystal of an antimicrobial substance for X-ray crystallography. (B) Chemical structure constructed by X-ray crystallography data. The structure is identified to Fluviol C (IUPAC name: 3-Methoxy-7-methyl-7H-pyrazolo[4,3-e][1,2,4]triazine). (C) Minimum inhibitory concentration (MIC) of Fluviol C against fish pathogens was measured.

### Effects of Fluviol C on Fish Pathogens and Fish Epidermal mucus bacterial community

To evaluate the antimicrobial activity of Fluviol C (FluC) against fish pathogens, we determined its minimum inhibitory concentration (MIC). FluC inhibited the growth of a range of Gram-negative and Gram-positive fish pathogens at concentrations between 0.5 and 32 µg/mL (Fig. 6C), consistent with the antimicrobial spectrum observed for the KH- ZF1 strain (Fig. S3).

We next evaluated the effect of FluC on the zebrafish epidermal mucus microbiota. Before administration, we determined a safe concentration range. Injured fish were transferred to flasks and maintained at the same water temperature as used in the infection experiments. FluC was added to the water at various concentrations, and fish survival was monitored (Fig. S9). While administration of KH-ZF1 cells at 10⁷ CFU/mL (OD₆₀₀ = 0.01) showed no toxicity to zebrafish, FluC displayed toxicity at concentrations exceeding 100 ng/mL (Fig. S10). Therefore, subsequent experiments were conducted using FluC at concentrations below 50 ng/mL (sub-MIC levels), and the epidermal bacterial communities of surviving fish were analyzed at one (12.5, 25, and 50 ng/mL) and two (50 ng/mL) days after administration.

At one day post-treatment, the relative abundance of the genus *Pseudomonas* increased compared to the untreated control (Fig. 7A), with the strongest effect observed in fish treated with 50 ng/mL FluC. By the second day, an increase in the abundance of *Flavobacterium* was also detected.

**Fig. 7.**
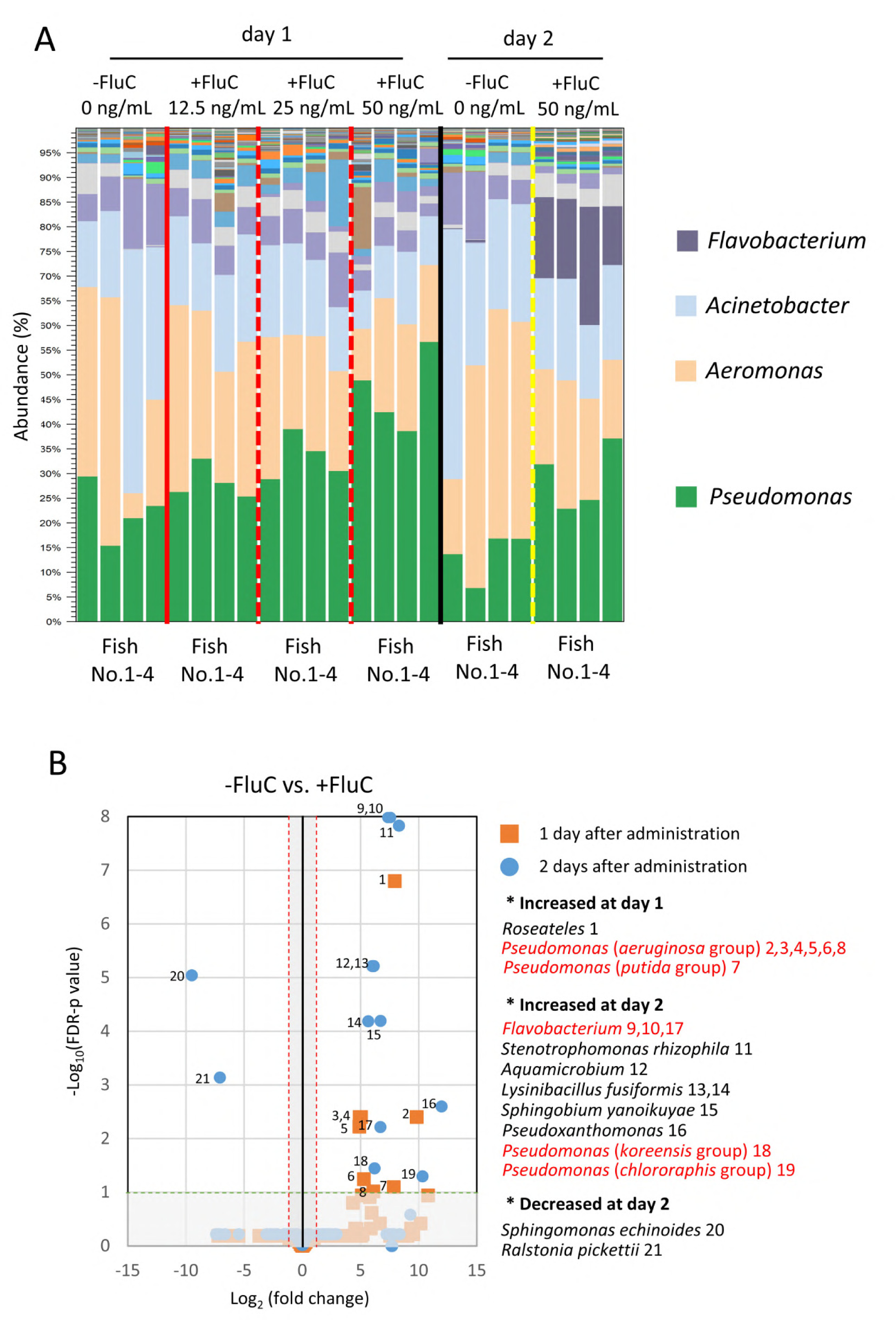
FluC promotes microbial substitution in epidermal mucus bacterial flora by administrating in rearing water. (A) Epidermal mucus bacterial flora analysis at 24 and 48 hours after administration of FluC. Final concentration of FluC in rearing water was 50 ng/mL. (B) OTUs significantly increased (No.1-19) or decreased (No. 20 and 21) at 1 and 2 days after FluC administration. Red gates represent above two-fold-increase or decrease against -FluC group and green gate represent FDR p-value under 0.1.

Differential abundance analysis of operational taxonomic units (OTUs) revealed significant increases in OTUs affiliated with dominant genera such as *Pseudomonas* (e.g., *aeruginosa* and *putida* groups) and minor genera including *Roseateles* one day after treatment. Two days after FluC exposure, OTUs associated with Flavobacterium, *Pseudomonas* (*koreensis* and *chlororaphis* groups), and several other genera were significantly altered (Fig. 7B). These findings demonstrated that sub-MIC levels of FluC can induce microbial substitution within the fish epidermal mucus microbiota.

## Discussions

Fish epidermal mucus functions not only as a physical barrier to environmental pathogens but also as a habitat for diverse microbial communities, including symbiotic and commensal bacteria (19–21). These communities are thought to contribute to host health and pathogen defense through complex interactions with invading microbes and the host immune system (26).

Epidermal injuries significantly increase the risk of pathogen entry and infection. Studies have shown that even minor abrasions can facilitate infection by fish pathogens such as *Flavobacterium psychrophilum* in ayu (*Plecoglossus altivelis*) (27), *Vibrio anguillarum* in zebrafish (28), and *Yersinia ruckeri*, as previously demonstrated (23). Such injuries are common in aquaculture due to high stocking densities and handling stress, highlighting the importance of strategies to protect the epidermis, including the use of beneficial microorganisms present in the mucus.

Several studies have isolated epidermal bacteria that inhibit fish pathogens. For example, isolates from brook and rainbow trout suppressed *F. psychrophilum* (16, 29). In our previous work, we identified antimicrobial *Pseudomonas* spp. in rainbow trout epidermal mucus (22). In this study, we identified *Pseudomonas mosselii* strain KH-ZF1 from zebrafish mucus (Table 1, Fig. 1), reinforcing the view that beneficial bacteria commonly inhabit fish skin. However, their utility for infection prevention remains uncertain, as prior studies report inconsistent outcomes: some found no protection (30), others observed reduced mortality (29). Our data suggest that timing and frequency of administration critically influence protective effects (Fig. 4A).

To clarify these discrepancies, we examined microbial community changes in epidermal mucus, gills, and gut after KH-ZF1 administration. KH-ZF1 transiently dominated the mucus microbiota and induced “microbial substitution” (Fig. 3C, Fig. S5, Fig. 4C). Similar dynamics were previously observed in trout mucus with antimicrobial *Pseudomonas* strains (22) and under stress or antibiotic exposure (23), suggesting antimicrobial activity—whether from drugs or microbes—can restructure the skin microbiota.

While antibiotic-induced dysbiosis (31) often enhances pathogen colonization and infection (23), KH-ZF1-induced microbial substitution appeared protective. KH-ZF1 given immediately after infection led to *Y. ruckeri* dominance, but a follow-up dose on the next day allowed KH-ZF1 to dominate and suppress pathogen growth (Fig. 4B, 4C). These results indicate that sequential microbial perturbation enables KH-ZF1 to occupy ecological niches and block pathogen colonization. Understanding such temporal dynamics is essential for designing effective biocontrol strategies.

To explore the mechanism behind KH-ZF1’s effects, we identified its antimicrobial compound. KH-ZF1 was found to produce Fluviol C (FluC), also known as pseudoiodinin (Fig. 6A, 6B), a pyrazolotriazine previously described as a pigment from *Pseudomonas fluorescens* var. *pseudoiodinum* (24). A recent study also identified FluC from *P. mosselii* strain 923 (25). Since *P. mosselii* was reclassified as distinct from *P. fluorescens* in 2002 (32), *P. fluorescens* var. *pseudoiodinum* likely belongs to *P. mosselii*.

FluC was initially mischaracterized chemically, but the structure was corrected in 2006 (33) Our chemical analysis confirmed the identity of the compound (Fig. S7, Fig. S8, Table S1), consistent with updated reports (33). Although earlier studies reported high MIC values (5–200 mg/mL) (34), our data and recent findings (Fig. 6C) demonstrate that MICs for several pathogens range from 0.5 to 32 µg/mL, despite differences in the bacterial strains used. These results indicate that the compound possesses strong antimicrobial potency. In another study, the MICs of FluC derived from *P. mosselii* 923 against the plant pathogens *Xanthomonas* spp. and *Magnaporthe oryzae* were reported to be 0.5 and 8.25 µg/mL, respectively (25). The recent report, together with our result, suggest that FluC exhibits antimicrobial activity at much lower concentrations than previously reported.

Despite its potent antimicrobial effects, the mechanisms underlying FluC’s antimicrobial activity and its role in microbial community substitution remain unclear. Notably, even sub-MIC concentrations were sufficient to alter the skin microbiota (Fig. 7A), suggesting that FluC may exert effects beyond direct bactericidal action. Differential OTU analysis revealed an enrichment of *Pseudomonas* and *Flavobacterium* taxa following FluC exposure (Fig. 7B). The sub-MIC levels of FluC might enhance the activity of resident bacteria, potentially contributing to indirect suppression of pathogens. For practical use, understanding metabolite localization and production timing is critical. FluC is toxic to zebrafish even at sub-MIC levels, so uncontrolled production could be harmful. However, KH-ZF1 administration did not reduce fish survival (Fig. S10), suggesting in situ toxicity is limited or transient. KH-ZF1 aggregated at injury sites (Fig. 3B), likely concentrating FluC locally to block pathogen entry while minimizing systemic exposure. Its temporary adhesion to mucus (Fig. 3A) may further support its protective effect. Thus, localized and transient colonization likely underpin its efficacy. We attempted HPLC quantification of FluC in the epidermal mucus after KH-ZF1 administration. However, interfering compounds in the mucus matrix overlapped with the target peak, precluding accurate detection. Further examination might need for measuring concentration of FluC in the epidermal mucus.

In summary, our findings highlight epidermal microbiota modulation as a promising strategy for aquaculture disease control. Understanding the roles of beneficial microbes and their metabolites in host interactions can inform alternatives to antibiotics and support sustainable fish health management.

## Materials and Methods

### Isolation of bacteria with antimicrobial activity, bacterial strains used, and the maintenance of bacterial cell culture

To isolate bacteria from zebrafish epidermal mucus, mucus was collected from anesthetized fish using sterile cotton swabs and suspended in 1 mL ultrapure water. Aliquots (200 μL) were plated on NB2 agar (10 g/L peptone, 10 g/L beef extract, 5 g/L NaCl, 1.5% agar), enriched cytophaga agar, modified Zobell 2216E agar (0.8% NaCl), and TSA (211825, Becton and Dickinson, Franklin Lakes, NJ). Plates were incubated at 28°C for 2 days. Colonies were subcultured, streaked, and single colonies obtained. Antimicrobial activity was screened by cross-streaking (see "Detection of Antimicrobial Activity of KH-ZF1") and purification was repeated five times. Isolates were identified via full-length 16S rRNA gene sequencing; sequence alignment analysis and a neighbor-joining tree was constructed using used CLC Genomic Workbench 11.0.1 (QIAGEN, Venlo, Netherlands).

*Yersinia ruckeri* strains NVH 3758 and DSMZ18506 (from Dr. Dirk Linke, University of Oslo) and NVH3758::lacZ (22) were grown in LB (Miller) at 28°C with 115 rpm shaking for 24 h. *Pseudomonas mosselii* KH-ZF1 was maintained on NB2 agar and cultured in NB2 or TSB at 20°C. KH-ZF1::mCherry was generated via conjugation from *E. coli* S17-1 λ pir carrying pBSL118_23119-mCherry, constructed by ligating the J23119 promoter and mCherry gene into pBSL118 (35). Other strains are listed in Table S2.

### Animal experiments

Adult zebrafish (Danio rerio) were obtained from MASUKO Co., Ltd. and maintained in 60 × 30 × 36 cm aquaria for at least 2 weeks. Fish were fed Tetra Min Super 17653 (Spectrum Brands Japan) every 12 h using a Tetra Auto Feeder AF-3, and water temperature was kept at 28°C using a SAFE COVER HEAT NAVI SH80 (GEX, Osaka, Japan).

For KH-ZF1 administration, seven or eight fish were injured as previously described (23), placed in a 500 mL flask with 300 mL sterile water at 20°C with aeration, and treated with 1 mL aliquots of KH-ZF1::mCherry (OD_600_ = 1.0) multiple times. After 24 h, fish were euthanized and skin, gills, and gut were sampled. For infection prevention, fish were injured, held for 24 h at 20°C, challenged with *Y. ruckeri*::lacZ for 6 hours as previously described (23), and followed by treated with KH-ZF1 (Fig. S6). Survival, CFU counts from epidermal mucus (collected by vortexing fish in 5 mL sterile water for 1 min), and bacterial flora analysis were performed. For CFU count of KH-ZF1::mCherry, *Pseudomonas* isolation agar (292710: Becton and Dickinson) with 50 μg/mL kanamycin was used for selection medium. For counting *Y. ruckeri*::lacZ, LB medium with 20 μg/mL X-gal and 50 μg/ml kanamycin was used. For FluC administration, fish were injured and water temperature was changed similarly, and exposed to varying FluC concentrations (Fig.S10).

### Sequencing of 16S rRNA gene amplicon libraries

DNA was extracted using the NucleoSpin Tissue kit (Takara Bio, Otsu, Japan) following the protocol for difficult-to-lyse bacteria, as previously described (23). During amplicon library preparation, a negative control (without environmental DNA) and a positive control (containing known bacterial genomic DNA) were included. No amplification was observed in the negative control, while the expected bacterial sequences were successfully detected in the positive control amplicon library by using primers for V1-V2 or V3-V4.

For analysis of bacterial community in animal experiments, 16S rRNA gene amplicon libraries (V1-V2 region) were sequenced using iSeq 100 (Illumina, San Diego, CA) (2 3). The V1-V2 region was selected to increase identification of *Pseudomonas* species (3 6). The V3-V4 region was used for detection of the isolated bacterial species in the isola tion sources. For sequencing of V3-V4 region of 16S rRNA gene, PCR products from bacterial isolation sources were sequenced by Seibutugiken Co., Ltd., Japan by using Illu mina MiSeq (Paired-end reads, 2×300 bp). The sequences of the PCR primers for V3-V 4 amplification are as follows: 1st 341f, ACACTCTTTCCCTACACGACGCTCTTCCG ATCTCCTACGGGNGGCWGCAG and 1st 805r, GTGACTGGAGTTCAGACGTGTG CTCTTCCGATCTGACTACHVGGGTATCTAATCC. The data were used for ASV analysis by DATA2 (37) .

### *In silico* analysis of 16S rDNA data

Paired-end sequences of the 16S rRNA gene (V3–V4 region) were processed using the DADA2 pipeline (v1.30.0) in R (v4.2.3 or v4.3.2). After quality filtering (truncLen = c (290,250), maxEE = c(2,2), maxN = 0), error correction, and chimera removal, amplicon sequence variants (ASVs) were inferred. Taxonomic classification of each ASV was performed using the naïve Bayesian classifier against the DADA2-formatted SILVA refere nce database (version 138.2; https://benjjneb.github.io/dada2/training, accessed on 16 June 2025).

Representative ASV sequences were used for reference-based phylogenetic analysis. Sequence alignment analysis and a neighbor-joining tree was constructed using CLC genomic workbench 11.0.1 (QIAGEN) to investigate the phylogenetic relationships among major ASVs.

The relative abundance of bacterial taxa at the genus level was calculated from the ASV table (Supplementary data sets No.1). To visualize the bacterial community composition across samples, a stacked bar chart was created using Microsoft Excel. The most abundant genera (Top 30 genus) were displayed, while the remaining low-abundance taxa were grouped into a single category labeled as "Others."

For the analysis of V1-V2 region, only sense reads (∼150 bp) were used because adequate read numbers were not obtained after merging reads. Reads were trimmed and analyzed using CLC Genomic Workbench with the Microbial Genomics module as previously described (22). OTU clustering was performed according to the instructions provided by the software. The SILVA 16S rDNA database (version 132; https://www.arb-silva.de, accessed on 16 October 2021) was used as the reference for taxonomic assignment, with a similarity threshold of 99%. During this process, chimeric reads were removed. Reads that could not be classified at the 99% threshold were re-analyzed using a lower threshold of 94%. Reads showing less than 94% similarity to any reference sequence were discarded. OTU abundance tables were then generated (Supplementary Data Sets No.1, No.2 and No.3). Differential abundance analysis (Supplementary Data Set No.3) was performed using the OTU abundance tables derived from FluC-treated and untreated groups (Supplementary Data Set No.2 and No.3). A volcano plot was generated using the log2 fold-change and -log10 FDR p-values of each OTU with a maximum group mean greater than 15.

### Detection of antimicrobial activity

Cross-streak assays were conducted as previously described (22), by vertically streaking skin bacteria and horizontally streaking pathogens. Plates were incubated at 20°C for 1-2 days.

For co-culture, pre-cultured *Y. ruckeri* NVH3758::lacZ and KH-ZF1 were inoculated into NB2 (20 mL) to OD_600_ = 0.001. *Yersinia ruckeri* CFUs were measured on NB2 agar with 50 µg/mL kanamycin.

Disk diffusion assays were done using *Y. ruckeri* suspended in NB2 with 0.5% agar (OD_600_ = 0.01). Suspensions were poured over solid NB2 agar. Paper disks (49005010, ADVANTEC, 8 mm) were loaded with 40 μL of test samples (from 1 mL of medium concentrated via evaporator, resuspended in water). Plates were incubated overnight at 20°C. Zones of inhibition were recorded.

MICs were determined as described (38) by adding 2 μL of FluC to 98 μL of OD_600_ = 0.001 bacterial suspension in 96-well plates. Ampicillin (200 µg/mL) and hygromycin B (100 µg/mL) were controls. Media: NB2 for *Edwardsiella tarda*, *Vibrio ordalii*, *V. anguillarum*, *Bacillus subtilis*; TSB for *Streptococcus iniae*; YPAD for *Saccharomyces cerevisiae* and *Candida cylindracea*; MHB for others.

### Purification of antimicrobial substances

KH-ZF1 was pre-cultured in 30 mL TSB for 2 days, centrifuged (8000×g, 10 min), washed with M9 medium (3 times), and resuspended in 30 mL M9 + 0.4% glucose. Cultures were incubated at 20°C for 2 days. Supernatant was centrifuged (10,000×g, 10 min), filtered (SLHAR33SS, Merck), and pooled (400 mL).

Organic solvent extraction was then performed using 100 mL of supernatant, as shown in Fig. 5C. Silica gel chromatography used column (0152-03-10, Climbing Co., Ltd.) packed with cotton, sea sand (191-15955, Wako), and silica gel (44-60 µm, FUJIFILM Wako) in ethyl acetate. Elution used ethyl acetate/methanol gradients (10:0 to 5:5), followed by 100% methanol. Obtained all fractions were tested via disk diffusion.

UV-Vis spectra (Cary 60, Agilent) were used to determine HPLC detection wavelength. Active fractions obtained by silica gel chromatography were further purified by reverse-phase HPLC (LC-20AT, Shimadzu) on InertSustain C18 column (4.6 × 150 mm, 5 µm; GL Sciences Inc.) using 50% methanol, 0.5 mL/min, 40°C column oven, detection at 220 nm. Fractions eluting at ∼7.5 min were collected. From 100 mL of M9 medium, approximately 0.2-0.3 mg of the antimicrobial substance was obtained.

### Identification of antimicrobial substances

Molecular weight was determined by LC-MS (1200 Series, Agilent; Compact, Bruker) using ESI+ mode. 5 µL of purified fraction was analyzed under same HPLC conditions. NMR (Avance III 500, Bruker) was done on 0.8 mg sample in CDCl_3_ + 0.05% TMS (034- 17211, FUJIFILM Wako), using NES-600 tubes (Optima Inc.) at ∼4 cm height. ^1^H- and ^13^C-NMR (512 and 12,000 scans).

For crystallography, 0.8 mg sample in 300 μL chloroform was vapor-diffused against hexane in glass tubes (BC-MGT015, Bio Medical Science Inc.) in 50 mL sealed bottles for several days. Crystals were visualized under polarized light, and structure was analyzed using XtaLAB P200 (Rigaku), solved with CrystalStructure 4.2.2, and refined with SHELXL v2016/4 (39). Structures were validated with checkCIF (https://checkcif.iucr.org/).

## Author Contributions

HN and KH designed the experiments, and HN performed the isolation of bacteria and animal experiments. HN and KH wrote the manuscript. NS performed isolation and identification of FluC.

## Acknowledgement

The authors would like to thank Professor Dirk Linke (University of Oslo, Norway) for kindly providing *Yersinia ruckeri* NVH 3758, and Professor Yutaka Tamaru (Mie University) for supplying zebrafish for the initial screening. We are also grateful to Dr. Atsuo Suzuki and Dr. Shuhei Ohmura (Nagoya University) for their guidance in X-ray crystallographic analysis. We thank Mr. Yuya Tsukamoto for assistance in isolating the antimicrobial bacterium. We also extend our appreciation to Dr. Michio Homma of our laboratory for his critical reading and English editing of the manuscript.

## Data availability

The data presented in this study are openly available in the DDBJ Sequenced Read Archive (DRA) under the accession numbers DRR576011-DRR576031. X-ray crystallographic data can be provided if demanded to authors

## Supplemental materials

Supplementary information associated with this article can be found in the online version of the publisher’s website.

